# Data Science Issues in Understanding Protein-RNA Interactions

**DOI:** 10.1101/208124

**Authors:** Anob M. Chakrabarti, Nejc Haberman, Arne Praznik, Nicholas M. Luscombe, Jernej Ule

## Abstract

An interplay of experimental and computational methods is required to achieve a comprehensive understanding of protein-RNA interactions. Crosslinking and immunoprecipitation (CLIP) identifies endogenous interactions by sequencing RNA fragments that co-purify with a selected RBP under stringent conditions. Here we focus on approaches for the analysis of resulting data and appraise the methods for peak calling, visualisation, analysis and computational modelling of protein-RNA binding sites. We advocate a combined assessment of cDNA complexity and specificity for data quality control. Moreover, we demonstrate the value of analysing sequence motif enrichment in peaks assigned from CLIP data, and of visualising RNA maps, which examine the positional distribution of peaks around regulated landmarks in transcripts. We use these to assess how variations in CLIP data quality, and in different peak calling methods, affect the insights into regulatory mechanisms. We conclude by discussing future opportunities for the computational analysis of protein-RNA interaction experiments.

## Introduction

RNA binding proteins (RBPs) are key orchestrators of post-transcriptional RNA regulation. They determine the fate of a transcript throughout its life-cycle; directing regulatory stages including splicing, polyadenylation, localisation, translation, stability and degradation. Over a thousand human RBPs have been annotated and identified by mass spectroscopy studies (1, 2). RBPs specify their RNA binding sites by recognising a combination of features, including RNA sequence motifs, RNA modifications, RNA structural motifs, and interactions with additional RBPs that bind at nearby loci (3). Each transcript interacts with many different RBPs to assemble into a ribonucleoprotein complex (RNP), which changes as the RNA passes through the various regulatory stages. RNP formation depends on the abundance of RNAs and RBPs in each cell type and on the post-translational modifications of these RBPs, and is sensitive to the competition between multiple factors for overlapping binding sites (4).

Due to the combinatorial and dynamic assembly of RNPs, it is crucial to identify the protein-RNA interactions that form within cells. Crosslinking between RNAs and proteins can be achieved by UV-C irradiation at 254 nm due to the photoreactivity of RNA bases (5). This has been exploited by “UV **c**ross**l**inking and **i**mmuno**p**recipitation” (CLIP) method that relies on UV light to crosslink covalently proteins to RNAs in intact cells or tissues, followed by purification and sequencing of RNA fragments that were crosslinked to an RBP-of-interest (6). Over the last 15 years, many variant protocols of CLIP have been developed, and in combination with high-throughput sequencing, they have led to a wealth of data, encapsulating transcriptome-wide binding profiles of hundreds of RBPs in multiple species, tissues and cell lines (7). The original CLIP and the derived variants all rely on sequencing, therefore we use the term ‘CLIP’ to refer generically to protocols that purify covalently crosslinked protein-RNA complexes and then sequence the bound RNA fragments. In contrast, we use the term CLIP-seq to refer to the protocol that was used by the first publication employing this term, which relied on readthrough cDNAs for data analysis (8) (Table 1, Supplementary Table 1).

Two orthogonal approaches to the analysis of CLIP data differ by their focus either on a specific RBP, or on the interacting transcripts. The RBP-centric approach aims to identify the RNA sequence, structure and other features that are in common across the binding sites of an RBP across the transcriptome, in order to unravel the mechanisms underlying the specificity of these interactions. This approach also aims to identify functional relationships between the bound RNAs, and their common regulatory principles. For example, the earliest CLIP studies of Nova proteins demonstrated that most RNA targets encode proteins with synaptic functions, identified the features of clustered YCAY motifs that are enriched at the endogenous binding sites, and defined the RNA map that demonstrated position-dependent activity of Nova at regulated exons and polyadenylation sites (6, 9, 10). The RNA-centric approach, on the other hand, examines the binding positions of a broad spectrum of RBPs on a specific transcript or sets of transcripts. This approach requires integration of CLIP datasets for multiple RBPs, which can be achieved by comparing available CLIP data that has been published by multiple research groups, and especially the eCLIP data produced as part of the ENCODE project (11). The results of both approaches need to be integrated with other methods to fully unravel the functions and mechanisms of action of RNPs (Box 1).

### Sidebar (Box 1)

#### RNA maps: integrating CLIP with orthogonal methods

An RNA map is a conceptually simple yet powerful tool that was initially developed to explore the functional impact of Nova binding motifs on splicing to predict Nova’s action genome-wide (9). It visualises the positional distribution of binding sites (commonly CLIP peaks or motifs) of the target RBP around ‘regulated landmarks’ in transcripts (such as alternative exons for splicing regulators). Landmarks are defined by an orthogonal method, for example by RNA-seq analysis of RBP knockout cells or tissues to identify the regulated exons. The distribution around each regulated landmark can be visualised as a heatmap, or summarised as a metaprofile (Figure 2d). To gain functional insight, the distribution around ‘control landmarks’ (such as unregulated exons) should also be plotted or used to determine binding enrichment, thus providing a sense of scale when comparing across experiments. The control variability can be examined using bootstrapping to determine the significance of enriched binding (97). To simplify implementation for general users, the rMAPS and expressRNA web servers have been designed to generate RNA maps using motifs or CLIP peaks around regulated exons and polyA sites (97, 98).

These maps are of great value not only in assessing RBP function, but also in validating CLIP experiments, since the enrichment of CLIP peaks around RNA features regulated by the same protein can serve as evidence of data specificity. The proportion of regulated RNAs with CLIP peaks at expected positions also provides insight into the sensitivity of data. Here, we use RNA maps to examine the sensitivity and specificity of CLIP peaks obtained by different CLIP methods and different peak calling tools or parameters (Figures 3 and 4).

**Table 1:**
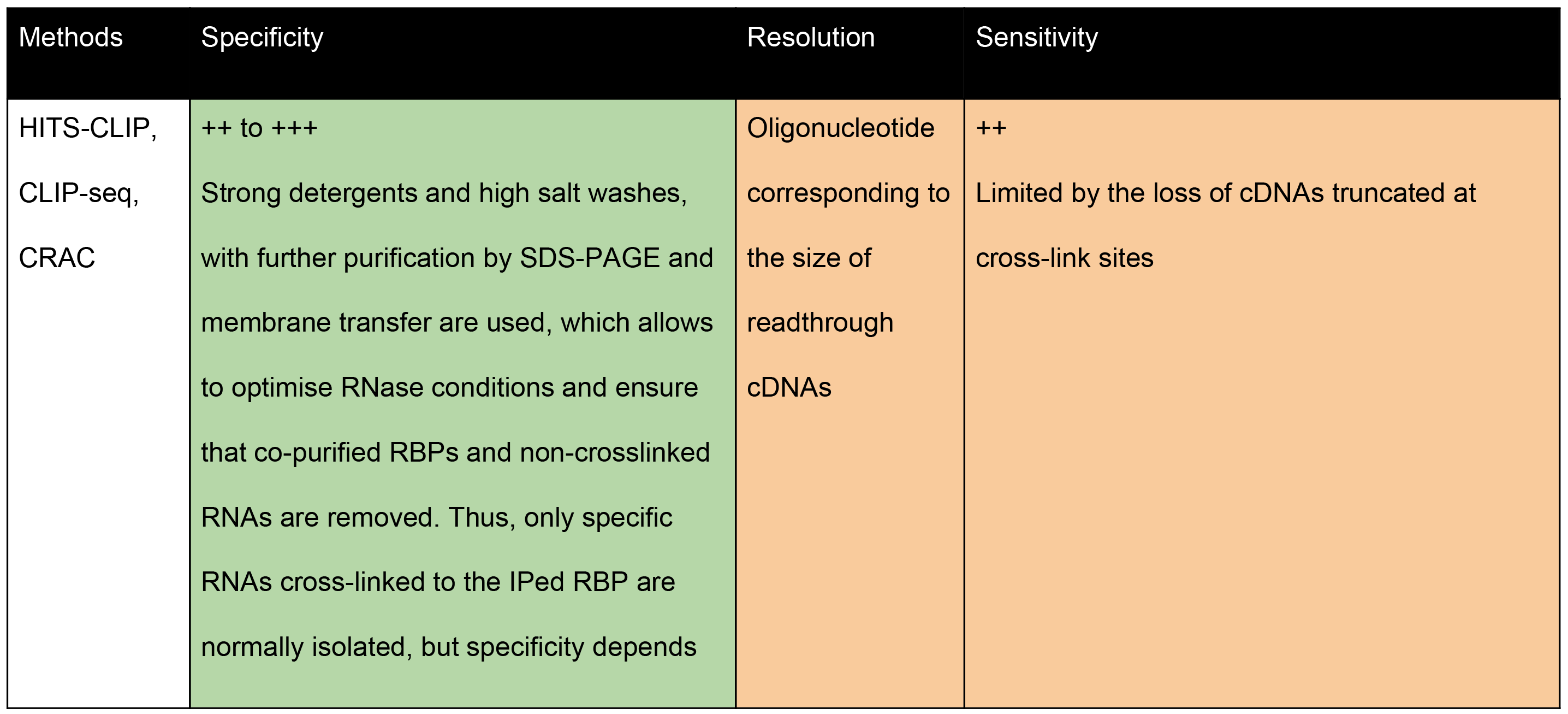
The central features of CLIP methods from the perspective of data analysis.

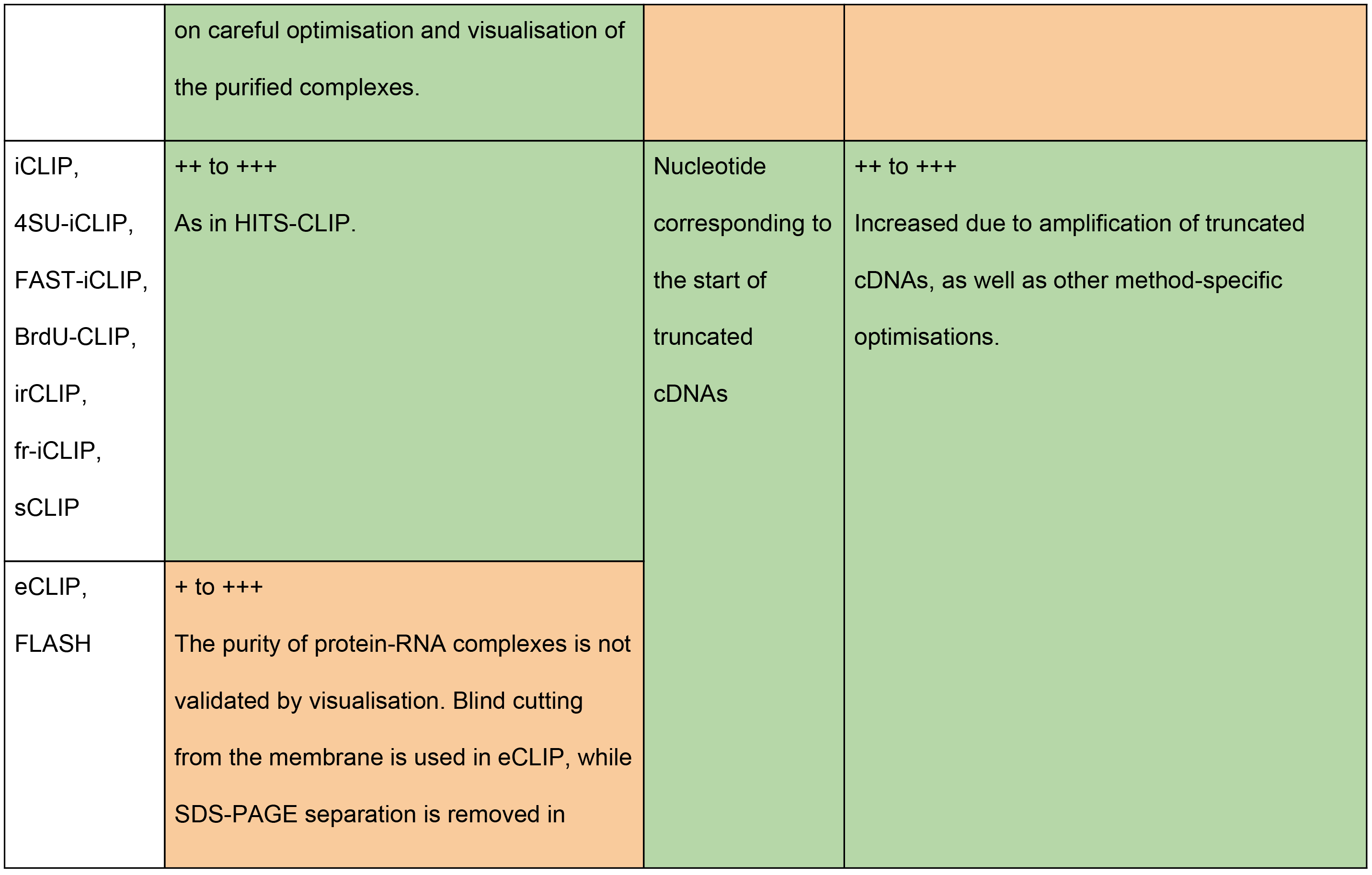

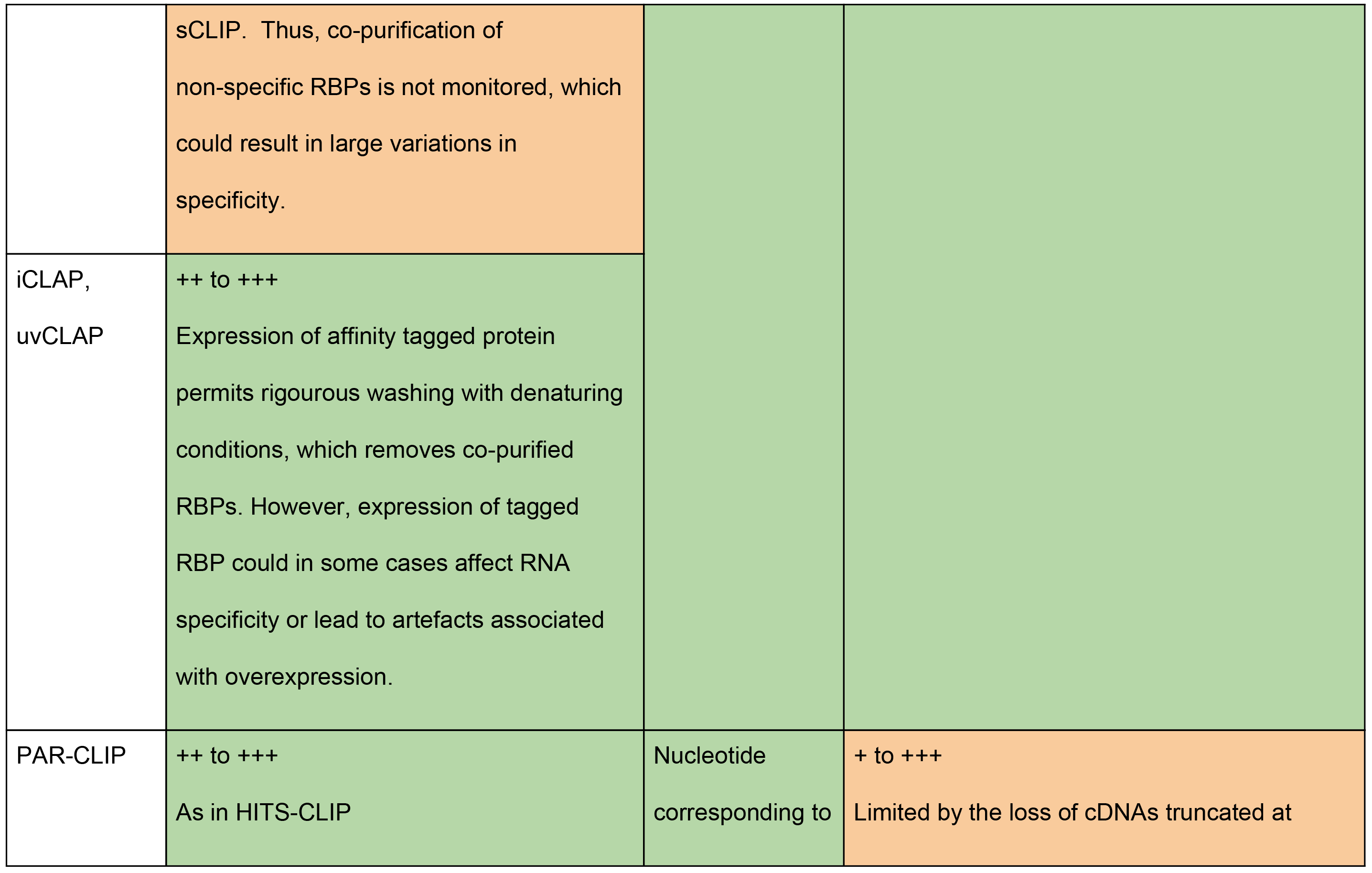

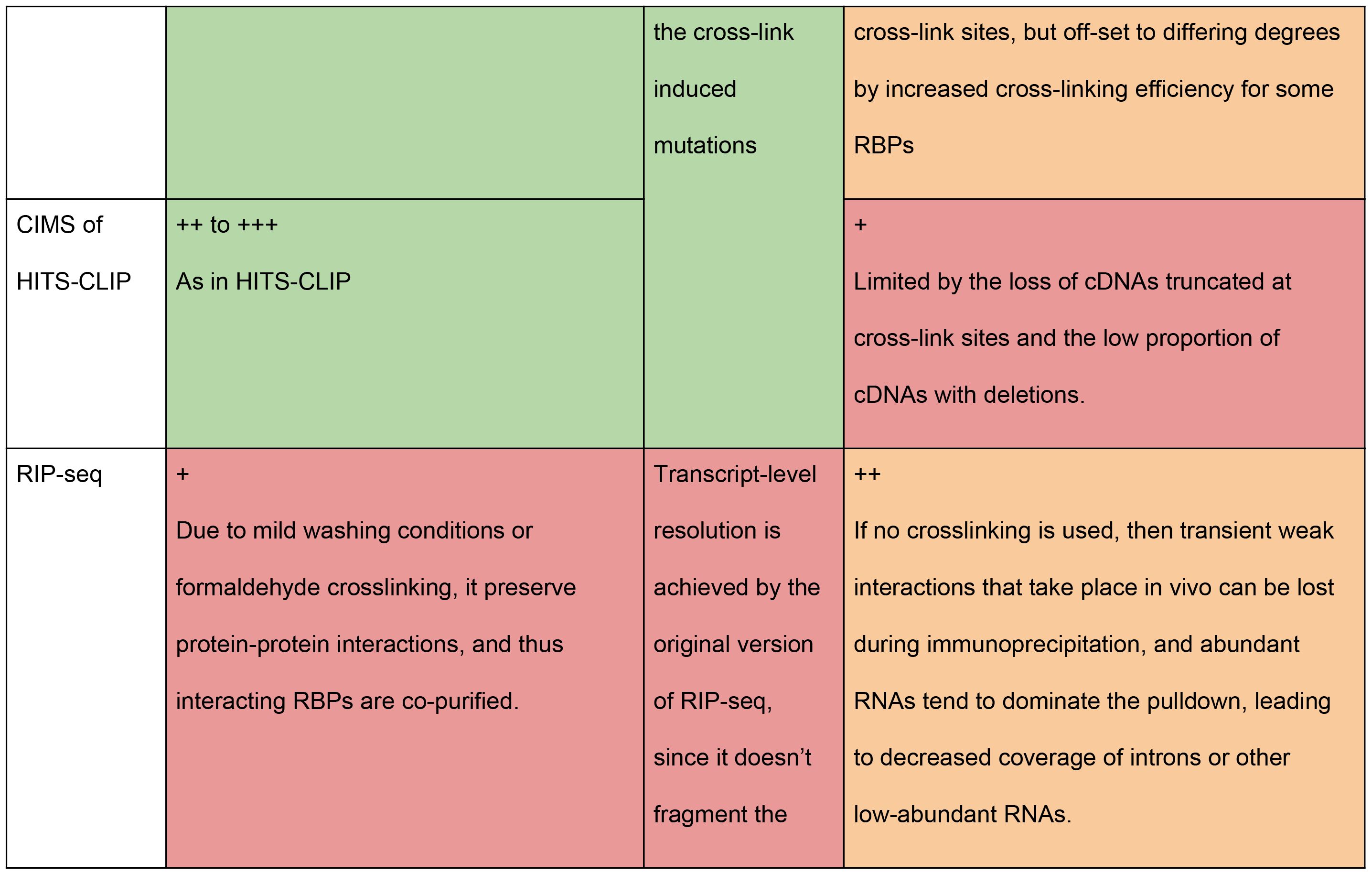

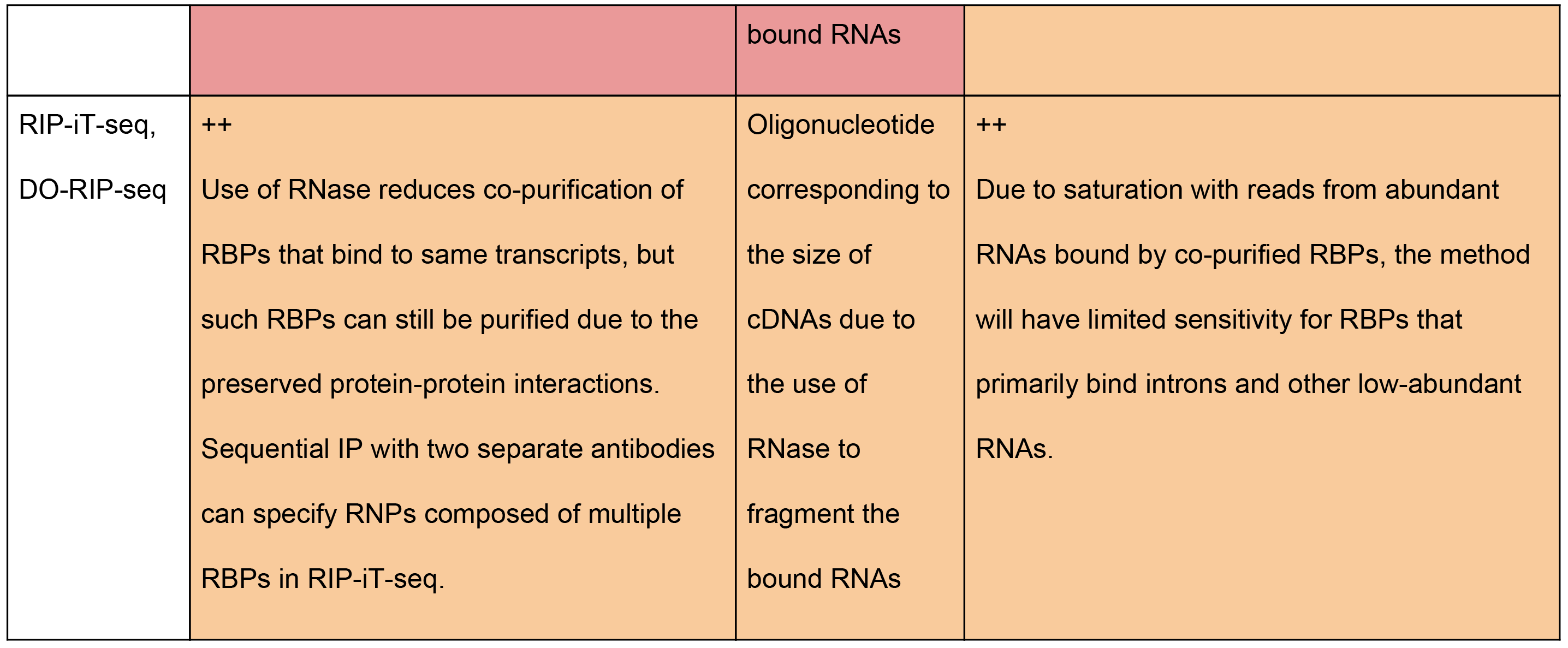

Table 2: Quality assessment of representative publically available CLIP data from different methods

Here, we review the computational and modelling methods, and use visualisation of enriched motifs and RNA maps to examine how the use of different methods impacts the biological insights gained from CLIP data. We start with a short evaluation of the experimental methods from a bioinformatic perspective; it is necessary to understand how the technical details of various CLIP protocols impact the specific requirements for the computational approaches. We then proceed through the primary stages of CLIP data analysis: i) quality control, ii) peak calling, iii) binding site modelling and iv) functional evaluation. At each stage, we explore the pertinent issues and potential pitfalls, through the lens of wanting to elucidate how RBPs recognise and act on specific transcripts. In the penultimate section, we broach what will likely become an important avenue of study in the near future: the integration of CLIP data from different RBPs. This will lay the foundations for developing an understanding of the complex network of RBP-RNA interactions. In concluding, we propose a set of standards as a framework for CLIP data analysis.

## Differences between CLIP methods from the perspective of data analysis

Despite the many variations of CLIP, its core principles mostly remain the same (7) (Figure 1). The covalent bond that is formed upon UV crosslinking allows the RNAs to be fragmented by a limited concentration of RNase after lysis, which is followed by purification of the RBP-of-interest under stringent conditions. Usually, an antibody is used to immunoprecipitate a specific RBP, which is separated on SDS-PAGE and visualised in complex with the crosslinked RNA fragments. The complex is then excised from the membrane and treated with proteinase K to remove the bulk of the RBP, leaving behind a short polypeptide at the crosslink site and releasing the RNA fragments. The fragments are then reverse transcribed into cDNAs, and the resulting cDNA library is sequenced. Initially, CLIP relied on traditional Sanger sequencing to identify 340 RNA fragments that provided the first glimpse into the binding sites of the neuron-specific Nova proteins (6), but now high-throughput sequencing enables us to gain a more comprehensive view across the transcriptome.

**Figure 1:**
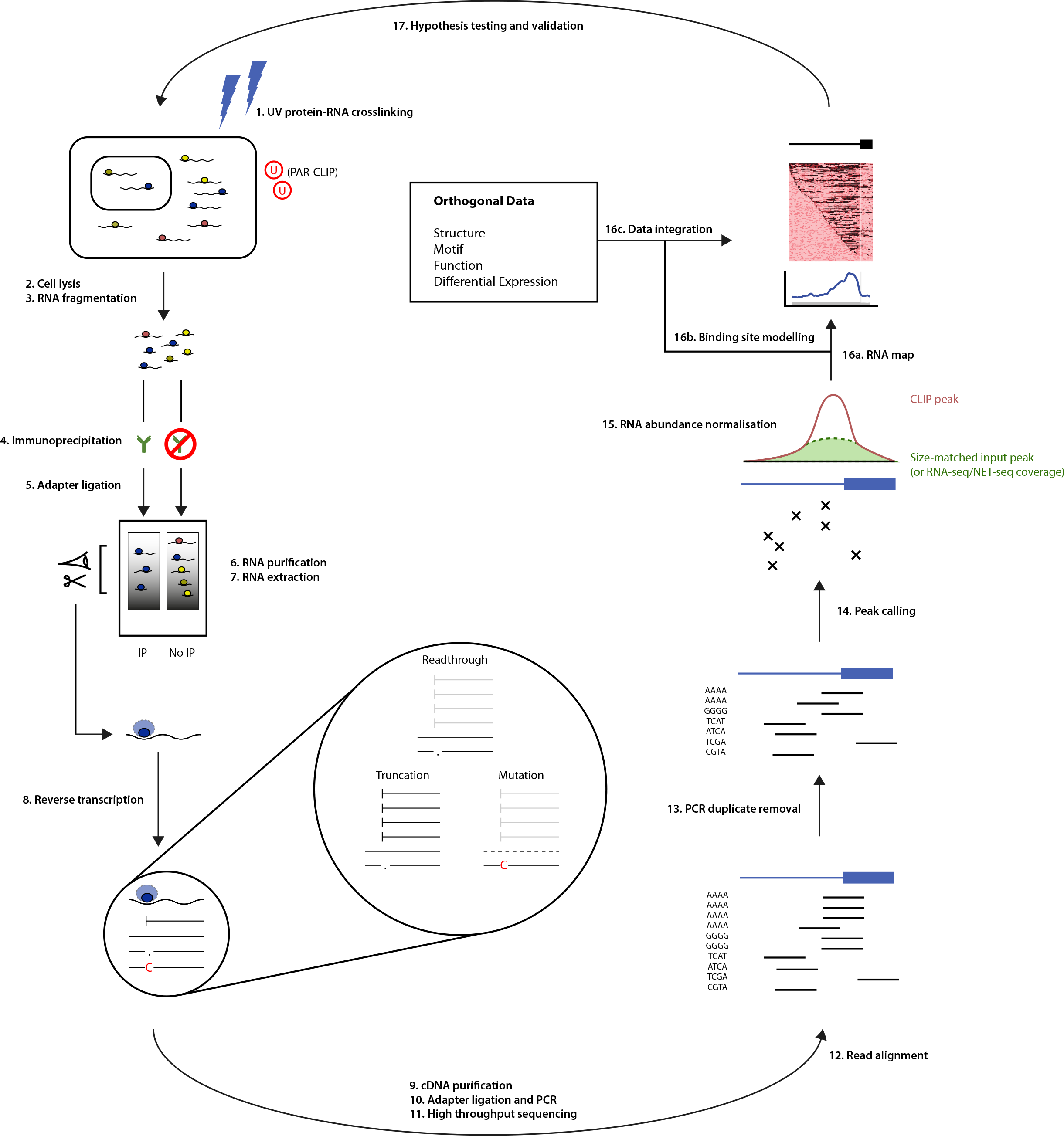
A computational biologist’s overview of the CLIP method. An outline of the key experimental (left) and computational (right) steps of the CLIP method. The experimental steps, common across most methods, are numbered according to (7) . Highlighted in the centre are the three primary data analysis approaches that rely on cDNA readthrough, mutation or truncation, depending on the type of CLIP protocol that was used to generate the data (related to Table 1). The cDNAs that are captured by representative protocols are marked in black, while those that are lost during reverse transcription in grey, and those that are discarded during analysis in dashed lines.

### Resolution and sensitivity

From the perspective of data analysis, CLIP methods can be divided into three principal approaches (Table 1, Figure 1). The division relates to the effect on reverse transcription of the polypeptide that remains at the crosslink site of fragmented RNAs. This can result in cDNAs that either: i) readthrough the peptide without any mutations; ii) readthrough the peptide but introduce a mutation at the crosslink site; or iii) truncate at the crosslink site.

The original CLIP method can only amplify cDNAs that fall into the first two categories, because both adapters that are required for cDNA amplification are ligated to the RNA fragments, and therefore the whole fragment needs to be reverse transcribed along with its adapters. This method employs UV-C light (254 nm) for crosslinking, which normally leads to only a minor proportion of cDNAs containing crosslink induced mutations (12). Therefore, binding sites for CLIP and its derived methods such as HITS-CLIP are usually assigned on the basis of the whole sequenced read (6, 10). Nevertheless, mutations, and especially deletions in CLIP cDNAs can help to increase the resolution of the method (13).

In PAR-CLIP, cells are pre-incubated with photoreactive ribonucleosides (usually 4-thiouridine, 4SU), which enables the use of UV-A light (365 nm) for crosslinking (14). Similar to CLIP, PAR-CLIP only amplifies cDNAs that fall into the first two categories, but it increases the proportion of cDNAs with mutations. About 50% of PAR-CLIP cDNAs normally contain thymidine to cytidine transitions at the crosslink site, which is the basis for binding site assignment by most tools developed for PAR-CLIP analysis (14). However, a large proportion of PAR-CLIP cDNAs lacks transitions, and longer cDNAs may contain more than one transition, and thus only a subset of cDNAs can be used for nucleotide-resolution studies (14).

iCLIP was developed to capture the third category of cDNAs that truncate at the crosslink site, in addition to the first two categories. This is achieved by ligating the second adapter to the cDNAs rather than the RNA fragments (15). In truncated cDNAs, the adapter is ligated exactly at the positions of their truncation. It has been estimated that approximately 90% of cDNAs in iCLIP truncate at the crosslink site (12, 16). Therefore, the nucleotide in the genome immediately 5’ of the aligned iCLIP cDNAs most often corresponds to the crosslink site. The same data analysis method applies to iCLIP and its more recent variants that also amplify truncated cDNAs, including irCLIP and eCLIP (17, 18).

The sensitivity of all CLIP methods is driven to a large extent by this choice between the three principal approaches. The relative crosslinking efficiency with UV-C or UV-A differs between RBPs (19), and this affects the relative sensitivity of CLIP versus PAR-CLIP methods. Both CLIP and PAR-CLIP lead to the loss of cDNAs truncating at crosslink sites, which in most cases represent ~90% of the total; therefore it is expected that iCLIP increases the sensitivity by a factor of ten. If UV-A crosslinking upon 4SU preincubation is beneficial, it can be combined with iCLIP in the variant termed 4SU-iCLIP (16, 20). As evident from the distribution of raw crosslink sites determined for PTBP1 by the different methods around its regulated exons, sensitivity can vary greatly between currently available data, with ~18% of repressed exons containing an iCLIP crosslink site at the peak position at 3’ splice site, compared to 4% containing an eCLIP crosslink (Figure 3a). This difference is not due to the choice of exons, since the sensitivity does not increase when using different eCLIP data or exons defined by RNA-seq analysis of knockout cells (Supplementary Figure 1).

### Specificity

The specificity of CLIP depends less on the choice of the three principal methods, and more on the stringency and validation of the steps required for purification of the protein-RNA complex of interest. Many RBPs participate in stable RNPs that do not dissociate even under the relatively stringent immunoprecipitation conditions of the standard CLIP, especially in the presence of RNA fragments that could help to stabilise them. Co-purified RBPs can have different RNA specificities and functions from the RBP-of-interest, and therefore ideally no additional RBPs should be co-purified to ensure high specificity. A denaturing condition is used by some CLIP variants to disrupt interactions with co-purified RBPs, but this is not possible when using antibodies that recognise the natively folded state of endogenous RBPs.

Separation of complexes by SDS-PAGE and membrane transfer, followed by their visualisation, along with the use of appropriate negative controls, is thus a crucial quality control step for methods that omit a denaturation step. Greater care needs to be taken when analysing data from methods that neither denature, nor visualise the complexes, such as eCLIP (17), since it cannot be assumed that the sequenced reads represent only RNAs in contact with the protein-of-interest. In these cases, careful computational quality control analyses, for example with the use of orthogonal data and RNA maps, should be used to examine specificity on a protein-by-protein basis. As evident by the analysis of PTBP1 RNA splicing maps (Figure 3b), the specificity of iCLIP is highest, since the silenced exons are specific in the enrichment at 3’ splice site, while enhanced exons contain enrichment downstream of the exons. The specificity is also high for eCLIP in spite of the low sensitivity, with enrichment at silenced, but not enhanced exons. However, specificity is low for irCLIP, where the enrichment at 3’ splice site for silenced exons is only slightly larger than the enhanced exons.

The potential for non-specific signal is higher for RBPs with low abundance or poor crosslinking efficiency. Crosslinking between RNAs and proteins requires close contacts between an amino acid and the nucleobase. Moreover, analyses of diverse CLIP datasets indicate that crosslinking efficiency of uridines and uridine-rich motifs is highest (12, 16), and therefore RBPs that contain such motifs in their binding site are expected to crosslink best - with such RBPs, especially if they are abundant, non-specific signal is not expected to be a major concern. However, low-abundant or poorly crosslinking RBPs, which likely includes many non-canonical or double-stranded RNA binding proteins, such as STAU1, may require denaturing conditions to ensure that the isolated RNA fragments are specific (21). Taken together, visualisation of protein-RNA complexes with SDS-PAGE analysis can validate the specificity of purification across a broad range of RBPs and conditions, thus simplifying downstream computational analyses.

## A considered CLIP analysis strategy

At its outset, CLIP data analysis follows a similar pipeline to most next-generation sequencing, but it diverges for experimental quality assessment and the subsequent determination and functional integration of the identified binding sites (Figure 1). We start by noting the nuances of read alignment that are particular to CLIP. We delve into the distinct analytical issues faced on account of the experimental choices, by detailing the CLIP quality measures that are necessary to appraise any results. We then consider the many challenges encountered in elucidating binding sites from the aligned reads. We end by looking at ways to distill the properties of these sites, and to relate them to biological functions.

### Read alignment

After standard quality assessment of the sequencing run, the pipeline turns to read alignment. Comprehensive benchmarking of RNA-seq read aligners has recently been undertaken and is outside the scope of this review (22). However, there are three factors that should be considered when tailoring this step to a CLIP experiment.

The first decision is whether to align to the transcriptome or the genome. The main advantage of transcriptomic over genomic alignment is increased sensitivity, with the proviso that only annotated mature transcripts are considered. However, for the majority of cases, where there is usually sufficient experimental sensitivity, alignment to the genome is preferred. This ensures appropriate assessment of the many RBPs that bind to pre-mRNA transcripts, in introns for example. Moreover, the use of a splice-aware aligner would accommodate those that bind to mature mRNA transcripts.

Second, the use of unique molecular identifiers (UMIs) in iCLIP and later methods accounts for the amplification biases introduced by PCR, but to be able to deconvolve UMIs for cDNAs that map to the same position, it is best to use uniquely aligned reads only. To maximise the fraction of reads that can be aligned uniquely, the originating cDNA needs to be sufficiently long. When cDNA lengths are greater than 35 nucleotides, high alignment rates can be achieved even to common RBP-bound repetitive elements, such as *Alus* (23, 24). However, if the RBP under study has a preference for other repetitive elements, such as microsatellite repeats or snoRNA (small nucleolar RNA) clusters, then a customised solution may be necessary. One option for these cases is to align reads to a consensus repetitive sequence (11). Another is to use expectation-maximisation to assign multi-mapped reads (25), but this approach means that mapping position cannot be used as part of the procedure to identify PCR duplicates.

Third, more technically, it is important to fine-tune some of the alignment parameters to the CLIP method that has been used. Care is required over the choice of number of mismatches allowed: too lax a setting will align reads with multiple sequencing errors. These may subsequently be identified spuriously as originating from different cDNAs when collapsing duplicates. It will also affect the sensitivity and specificity of the mutation-based methods. Specifically for the truncation-based methods, it is important to disable soft-clipping to ensure the crosslink site, reflected in the start of the read, is properly aligned.

### Quality control

Thorough quality assessment is imperative to understand the CLIP experiment, both for the appropriate assignment of binding sites and for integrating with other data sources. We propose that measures that evaluate cDNA complexity and specificity are explored in combination. We have applied basic quality assessments of these two measures to representative publically available data for the different variants of CLIP, which provides an estimate of the variation in existing data, and potentially helps set up standards for future experiments (Table 2, in preparation).

#### cDNA complexity

cDNA complexity informs on the sensitivity of the CLIP experiment. The total number of unique cDNAs gives an appreciation of the dynamic range of RBP-RNA interactions that can be detected. Complexity reflects a number of biological and technical factors: the abundance and crosslinking efficiency of the RBP, and the efficiency of immunoprecipitation, adapter ligation and cDNA library preparation. PCR duplication, although necessary for the method, can create difficulties for monitoring library complexity. Amplification of cDNA fragments is not uniform, but affected by sequence content and length. In the original CLIP protocols, it is therefore necessary to remove the duplicate reads, and consider only the unique reads to reliably count the cDNAs. The CLIP Tool Kit achieves this by collapsing the identical reads before alignment (26), but ideally reads are collapsed after alignment based on identical genomic start position, since this accounts for read variations that result from sequencing errors (13).

The current gold-standard is a more sophisticated approach that experimentally labels each cDNA as it is reverse transcribed (15, 27). This is done by introducing a UMI, which is a randomised sequence of nucleotides (hence also known as a random barcode or randomer), into the reverse transcription primer. After PCR amplification, the UMI remains as a hallmark of unique cDNAs. iCount and other tools developed for analysis of iCLIP data use UMIs in combination with the read start position to count the unique cDNAs accurately, and thus obtain reliable information about cDNA complexity and enable quantitative analysis of crosslinking at individual nucleotide positions. The use of UMIs is crucial to overcome the artefacts of PCR amplification and thus preserve the quantitative information present in the cDNA counts, which is particularly important in quantifying binding to high-affinity binding sites, and in abundant RNAs.

#### cDNA specificity

Establishing cDNA specificity is the most difficult evaluation, and yet the most important. It is mostly dictated by the purification of the RNA-RBP complex, hence the importance of optimising this process. Ground-truth is often not known, and the appropriate measurement may vary between RBPs, on account of differing sequence and structure specificities. In practice, often only circumspect or post-hoc approaches can be used.

The percentage of crosslink sites that occur in peaks is a basic measure of the capacity of the cDNA library to identify binding sites and gives some clue as to specificity (Table 2). Enrichment of RBP-specific, binding-related *k*-mers within peaks, compared to a suitable background region, also provides some reassurance. Enrichment of motifs (ascertained using alternative methods, such as RNAcompete (28)) within clusters of peaks gives another independent, complementary assessment of specificity. However, these last two will only work for RBPs that bind particular sequences or motifs; for those that do not, there will be little enrichment regardless. Finally, the integration of CLIP results with orthogonal data provides the best measure of specificity, but requires the availability of such data. RNA maps (detailed in Box 1) are an efficient approach for visualising crosslinking around transcriptomic landmarks that are relevant for the function of the RBP: exon-intron junctions of regulated exons, for example, for RBPs involved in splicing.

While overlapping cDNA starts are a measure of potentially high specificity of iCLIP data for crosslink clusters, they can also reflect the aforementioned sequence preferences of the UV crosslinking reaction, which needs to be taken into account. Moreover, overlapping cDNA ends in iCLIP (and both sides of cDNAs in HITS-CLIP and PAR-CLIP) reflect the preferences of the RNases used for fragmentation (29). The alignment of cDNA ends can lead to a generic misalignment of the starts of cDNAs of different lengths; while this was initially interpreted as possibly indicating presence of readthrough cDNAs (30), the RNase fragmentation biases were found to be the more likely cause (16). The alignment of cDNA ends correlates with an enrichment of /c-mers at cDNA ends, which is a useful tool to examine the biases introduced by RNA fragmentation. To avoid such biases, optimised RNase fragmentation conditions should be used to produce a broad range of cDNA lengths, which ensures that the full binding sites can be defined, which is particularly important for long binding sites (16). This also guarantees that the peaks identified by overlapping cDNA clusters are a true measure of data specificity, rather than an artefact of inappropriate RNase fragmentation.

### Peak calling

The main challenge of CLIP data analysis is related to the biological context of protein-RNA complexes. Binding cannot be classified into simple binary categories of specific and non-specific; instead RBPs bind RNAs with a range of affinities and kinetics (31). Some RBPs associate with RNA polymerase and transiently interact with many low-affinity sites on nascent transcripts before finding a high-affinity binding site, and others can spread over larger regions of RNA after finding a high-affinity sequence. Many assemble on RNAs combinatorially as part of larger complexes. While the probability that an RBP will crosslink repetitively to a clustered set of crosslink sites is generally increased at the sites with high affinity and favourable binding kinetics, the exact threshold for defining functionally relevant types of crosslink clusters depends on many factors, such as: the type of RBP under study, the type of bound RNA, the function under regulation, and the binding position relative to other regulatory complexes. Thus, there are no absolute thresholds that can be set to distinguish low-affinity, transient binding from high-affinity functional binding. This challenge could become insurmountable if data contain many non-specific sites that do not represent direct interactions of the specific RBP. It is thus of paramount importance to maximise the specificity of CLIP data experimentally, since this can ameliorate the computational analyses needed to identify the functionally relevant binding sites.

Peak calling is the first step towards identifying the RNA sites that are highly occupied by the RBP; those that are most likely of functional significance. The basic approach searches for the pile-up of aligned reads at specific positions on transcripts. In methods such as ChIP-seq and RIP, which tend to purify large protein-protein complexes as well as free DNA or RNA, the purpose of this step is largely to isolate the signal from the inherent background noise of the techniques. CLIP employs many unique experimental steps to remove such noise, including covalent crosslinking, RNA fragmentation, stringent purification and visualisation of purified protein-RNA complexes, and thus in a fully optimised experiment the mapped reads should almost exclusively correspond to the sites of direct protein-RNA contacts. Therefore, noise from non-specific background should not be a major concern for CLIP data analyses. As evidence of the high specificity of CLIP experiments, the raw whole reads from Nova (Figure 2c) and PTBP1 HITS-CLIP (Figure 3a), and the raw crosslink positions from PTBP1 iCLIP (Figure 3a), yield highly position-dependent enrichment on RNA splicing maps .

**Figure 2:**
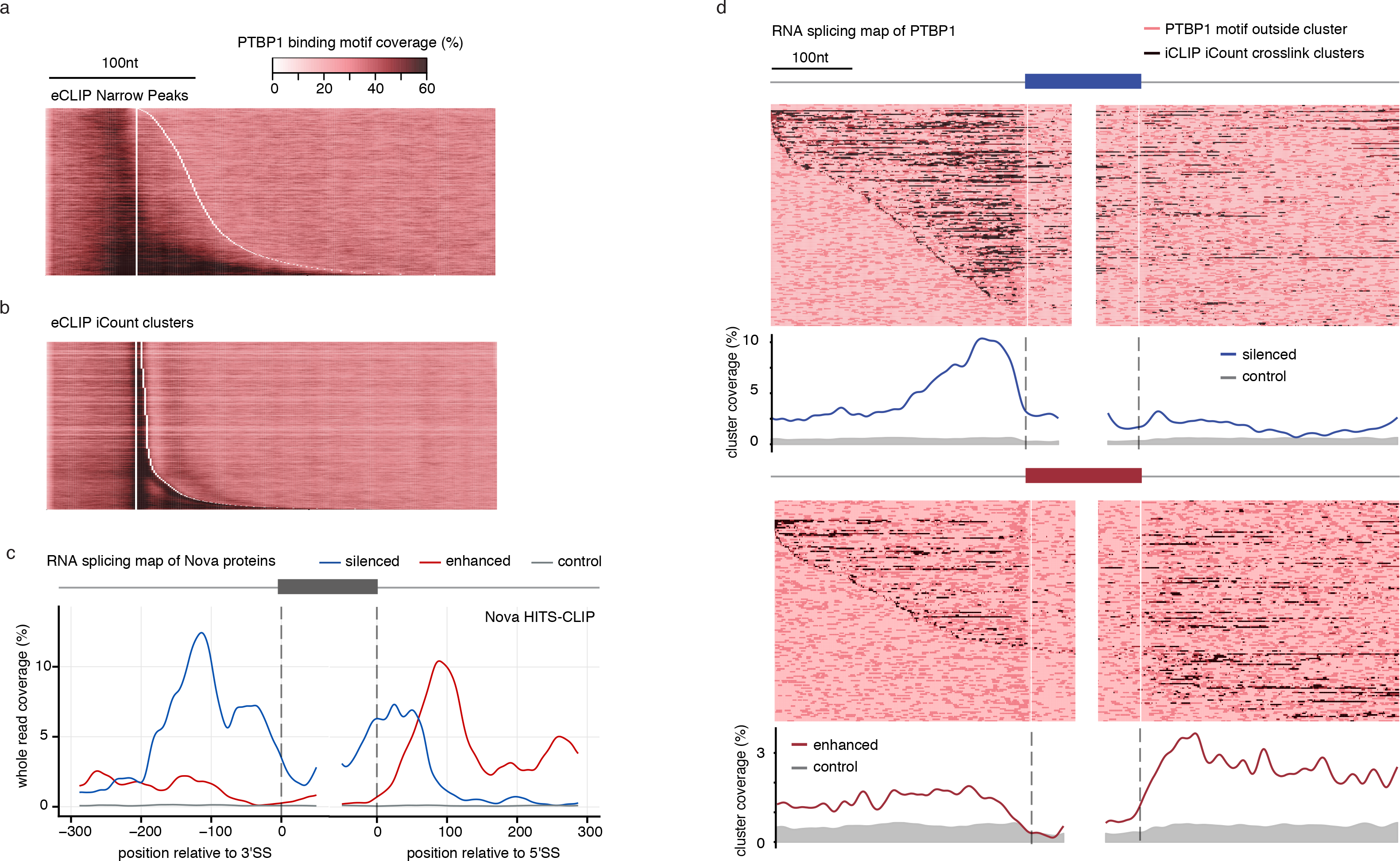
Visualisation of CLIP data: motif plots and RNA maps. a) The distribution of PTBP1 motifs from (16) are shown around eCLIP peaks that are defined as narrowPeaks and are available from the ENCODE website. This algorithm relies on the use of whole reads, which leads to misalignment of motifs and peaks, b) The iCount peak caller (15, 42) uses the starts of aligned reads to define the crosslink positions and peaks, which leads to good overlap with PTBP1 motifs, c) Integrating CLIP and orthogonal data allows further exploration of data quality using an RNA splicing map, which examines the distribution of clusters of assigned binding sites around repressed (blue) and enhanced (red) exons. This approach was first used with HITS-CLIP reads for NOVA in mouse brain (10). Here we assign a binding site to all positions in transcripts that overlap with at least one raw read, based on the 168,632 reads obtained by the original HITS-CLIP publication; even though we do not use peak calling, this results in high position-dependent enrichment that agrees well with the computationally predicted RNA map (9), thus highlighting the high specificity of raw CLIP data, d) RNA splicing map of PTBP1 iCLIP data from HeLa cells (16) is drawn in two ways with peaks called using iCount with 3 nucleotide clustering (15, 42). Regulated exons are defined using microarray data upon knockdown of PTBP1/PTBP2 in HeLa cells (99). Each row of the heatmap is a regulated exon with its flanking region; the positions of peaks are shaded dark; PTBP1 motifs inside or outside the clusters are shown as black or light red. The metaprofile of significant crosslink clusters is plotted below. The enrichment of peaks around regulated compared to control exons informs on the mechanisms of splicing regulation, and on the specificity of CLIP data. The code to reproduce this figure is available at https://github.com/jernejule/clip-data-science

**Figure 3:**
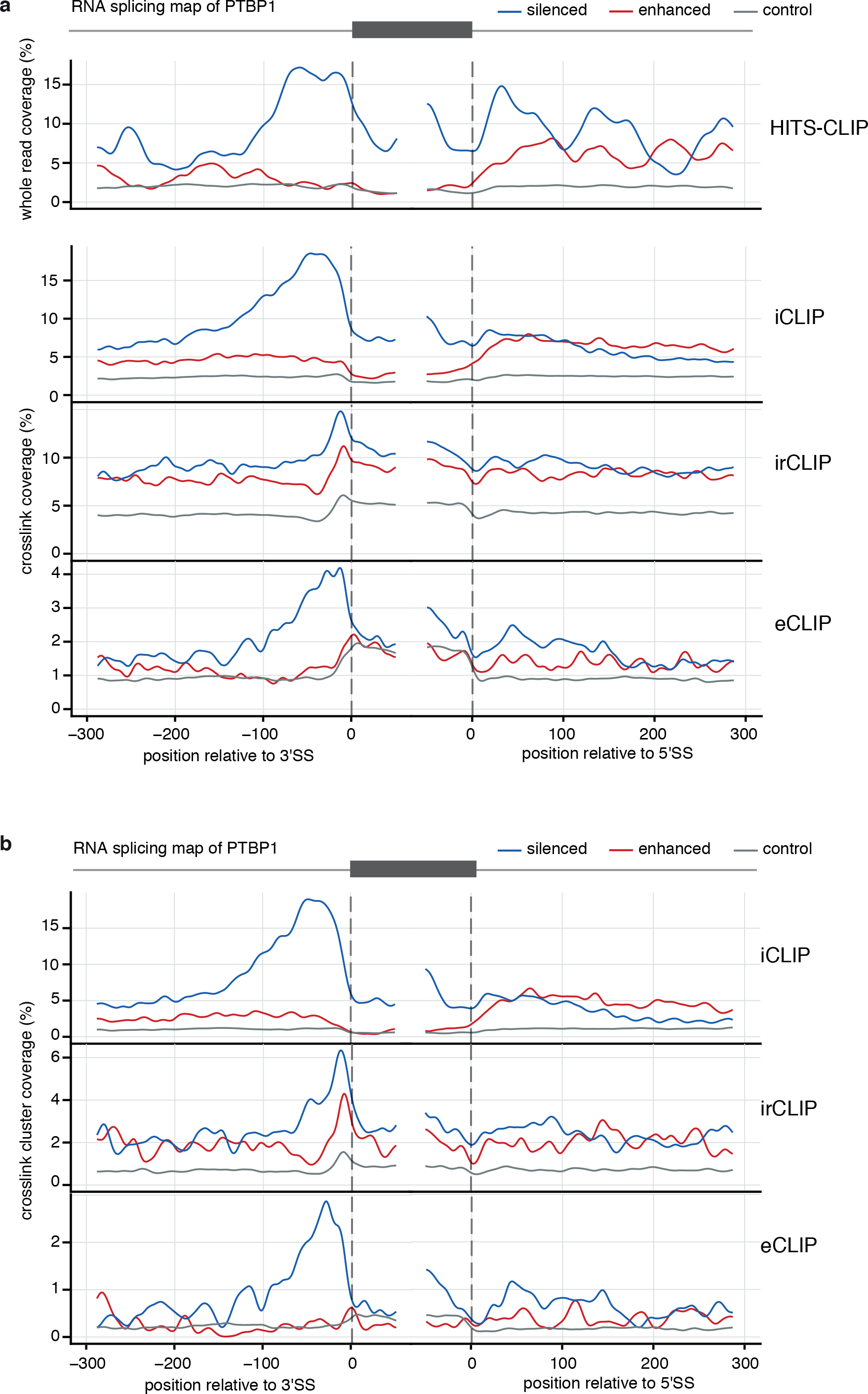
Using RNA maps to examine sensitivity and specificity of CLIP data. PTBP1 is an abundant RBP that crosslinks efficiently and follows position-dependent regulatory mechanism, and is thus a suitable RBP for data analysis via RNA map. The regulated exons were defined by analysis of splice junction microarray data with ASPIRE3 software (abs(dlrank)>1) upon knockdown of PTBP1/PTBP2 in HeLa cells (99). In a) we compare the raw data for different experimental methods, with whole reads from HITS-CLIP in HeLa cells (100),crosslink positions from irCLIP (18) and iCLIP (16) in HeLa cells, and eCLIP in HepG2 cells (11) . This demonstrates that CLIP data can lead to strong enrichments even without peak calling, but this depends on the specificity of data. In b) we analyse the effects of peak calling on the crosslink positions from different experiments, with data from irCLIP (18) and iCLIP (16) in HeLa cells, and eCLIP in HepG2 cells (11) all analysed using the iCount peak caller with 15 nucleotide clustering (15, 42). The code to reproduce this figure is available at https://github.com/jernejule/clip-data-science

Many peak calling tools have been developed (32), some specific for particular CLIP protocols, others more generally applicable (detailed in Supplementary Table 2). The large number of peak calling tools, which often come with adjustable parameters,may present a bewildering set of possibilities. This is further complicated by the different strategies in identifying the crosslink sites by the various experimental protocols (Table 1, Supplementary Table 1). Benchmarking of tools is challenging, owing to the differences in experimental protocols, and our limited understanding of the ground-truth regarding RNA binding sites in vivo (33, 34). Nevertheless, we attempt to demonstrate the impact of the different CLIP protocols and computational tools through use of the RNA maps, which combine CLIP with orthogonal functional data to derive an estimate of ground-truth from the perspective that RNA landmarks regulated by an RBP should contain its nearby RNA binding sites (Box 1). A comparison of peak calling by three tools demonstrate that all have similar specificity when using iCLIP data as input, with iCount leading to highest sensitivity, since it detects significant crosslink clusters at the peak position at 3’ splice sites of 25% of the repressed exons (Figure 4).

**Figure 4:**
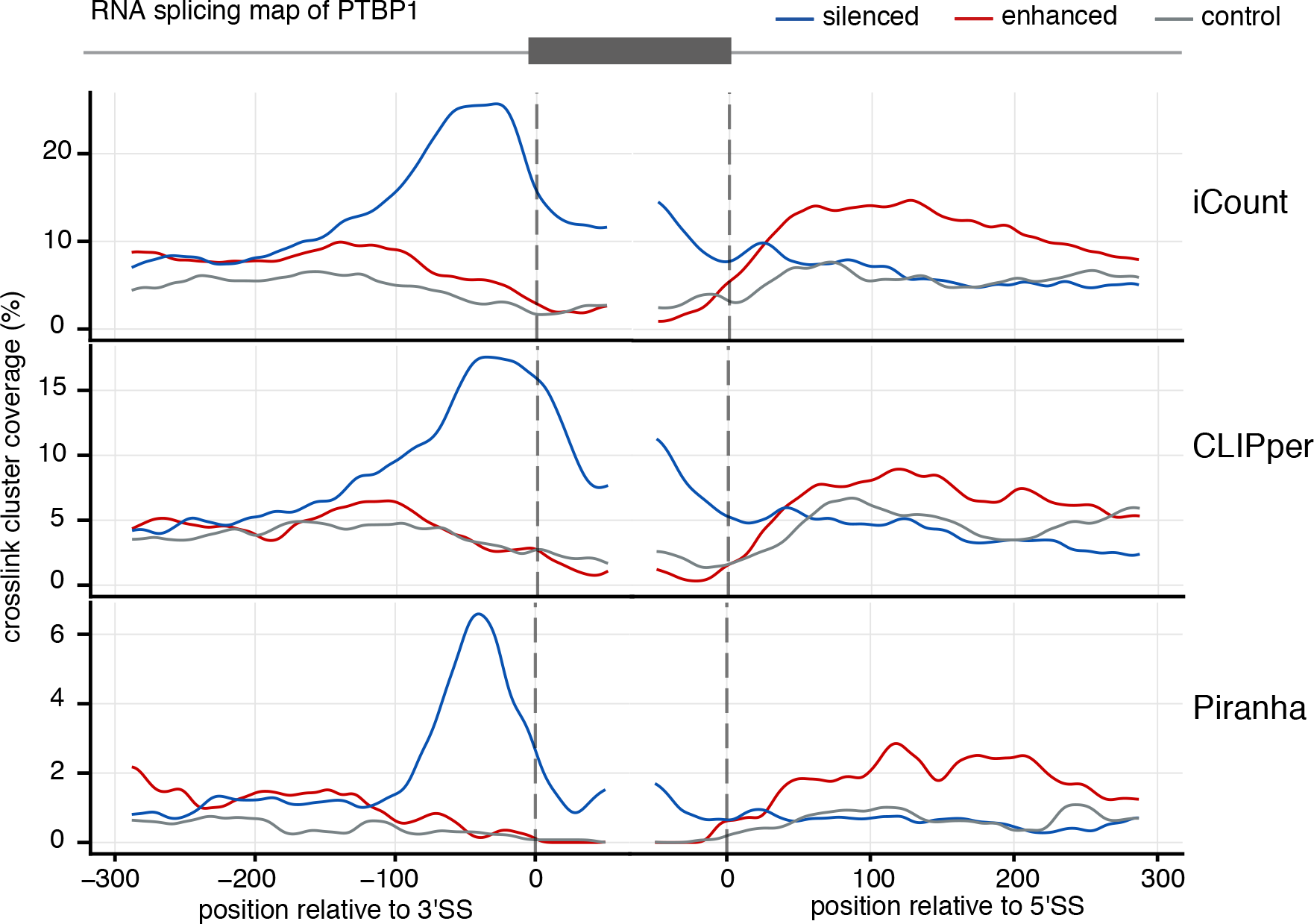
A comparison of different CLIP peak calling tools. RNA maps are used to demonstrate the differences in peak calling tools for the same iCLIP PTBP1 data set (16) . To demonstrate that the RNA maps can be reproduced by exons defined by a different data source, the regulated exons are defined using RNA-seq data following PTBP1 CRISPR knockout in K562 cells from the ENCODE website. We identified the skipped exons detected using rMATS (101) using junction counts only and a P-value threshold of 0.05 and FDR threshold of 0.1. Repressed and enhanced exons were defined using an inclusion level difference threshold of 0.05; control exons were selected as those with a P-value > 0.1, FDR > 0.1 and an inclusion level difference of < 0.001. We compare the peaks called using iCount (15, 42) (using a 15 nucleotide peak calling half-window and 30 nucleotide clustering window), Piranha (41) (using a 30 nucleotide bin size and 30 nucleotide merging window), and CLIPper (11, 44) (using default settings). For this dataset, Piranha and iCount have runtimes of ~2 minutes and ~7 hours respectively using 1 processor; CLIPper has a runtime of ~7 days using 20 processors. The code to reproduce this figure is available at https://aithub.com/ierneiule/clip-data-science

#### Challenge 1. What to use to call a peak?

The first consideration is how to use a read to define a peak. This differs for the mutation-based and the truncation-based CLIP methods. For mutation-based methods, it is important to distinguish a mutation from confounders, such as sequencing errors or single nucleotide polymorphisms or somatic mutations in cell lines. Early tools, such as PARalyzer, addressed this by setting a minimum number of mutations at a site, or by limiting the number of mismatches permitted during alignment (35). While a simple and effective way of reducing false positives, it has the disadvantage of also reducing the sensitivity of the experiment. To improve this, PIPE-CLIP models each event using a binomial distribution, with a success rate calculated from the read coverage (36). As a further refinement, wavClusteR uses a non-parametric, two-component mixture model to distinguish crosslink induced mutations from noise; it integrates this using a Bayesian network representation (37, 38).

For truncation-based methods, this seems more straightforward: the nucleotide upstream of the start of the read is the crosslink position (which we refer to as the ‘cDNA start’) and can be used to call peaks. However, there is a caveat. For the vast majority of the cDNAs, reverse transcription stops at the crosslink site, but it does still read-through at times (this provides the signal for the readthrough-based methods). In iCLIP experiments of most RBPs, however, over 90% of cDNAs terminate at the crosslink site (12). Therefore, as discussed, provided there are limited cDNA end constraints, and cDNA sizes cover a broad range of lengths, the use of the read starts assigns crosslink sites with no positional bias (16). Finally, 4SU-iCLIP uses 4SU for crosslinking as is done in PAR-CLIP, but then employs iCLIP protocol to prepare the cDNA library, which raises the question whether mutations (as in PAR-CLIP) or truncations (as in iCLIP) should be used. Analysis of PTBP1 binding motifs in 4SU-iCLIP cDNAs indicates that truncations report a more reliable estimate of crosslink sites than transitions (16). This still needs to be evaluated for additional RBPs.

It is important to use the appropriate marker to call peaks. The eCLIP ‘narrowPeaks’ publically available from the ENCODE consortium were defined using an algorithm that used whole reads. However, such use of whole reads leads to misalignment of binding sites and loss of resolution, as is evident from PTBP1 motif analysis (Figure 2a). This can be solved by the use of the truncation-based approach used by iCount algorithm that defines peaks based on the starts of mapped reads (Figure 2b), and a similar approach has been implemented also by the published eCLIP study (11). In summary, the use of whole reads is appropriate for the original variants of CLIP, mutations can be used as alternative sources for peak calling, such as T-to-C mutations in PAR-CLIP, while the read starts should be used for iCLIP and other methods that are optimised for amplification of truncated cDNAs.

#### Challenge 2. What is a peak?

The next problem is defining what constitutes a peak: how high and how wide does the pile-up of reads need to be? The former provides a guide as to the likelihood of a locus being a true binding site, while the latter considers when one binding site should actually be considered as two adjacent ones. This is of importance as some RBPs have narrow, focussed binding sites (e.g. PTBP1), whereas others bind more diffusely across a transcript (e.g. MATR3).

##### Peak height

The focus of most tools is calculating the probability of a binding site not belonging to a background CLIP read distribution (33, 39). Generally, a probability distribution is fitted to the count data; differences in the tools arise from the probability distribution function chosen to model the read counts and the generation of the background.

The majority of available tools use variations on a negative binomial distribution. This is often used for count data as it has the advantage of being able to account for overdispersion (i.e. if the variance of data is greater than the mean). ASPeak uses this distribution unmodified (40). Piranha (41) and PIPE-CLIP (36) use a zero-truncated negative binomial distribution. It has been shown for a range of RBPs and CLIP methods that this zero-truncated negative binomial distribution fits the count data better than simple negative binomial, or Poisson distributions (41).Piranha calculates the counts in user-defined bins across the genome; an appropriate size depends on the RBP. A zero-truncated negative binomial distribution is fitted to the data; bins where there is a higher read count than would be expected can then be selected as peaks, using a P-value threshold.

The iCount tool (15, 42), developed along with the iCLIP method, avoids fitting a specific distribution, but uses permutation analysis. The counts are randomly distributed a pre-defined number of times within a relevant region of interest (such as introns) on a gene-by-gene basis to generate a background. Then, the comparison of the observed distribution with the random one yields a false discovery rate. The primary disadvantage of this method is that, in order to generate meaningful random distributions, an annotation is needed to provide the regions of interest. A similar approach is used by the CLIP Tool Kit (26) and Pyicoclip (43).

CLIPper (44), the tool of choice of the ENCODE consortium (11), combines ideas from both these approaches. A false discovery rate, similar to iCount, is calculated in a first pass. However, by default the reads are randomly distributed within the entire gene, rather than a more localised region of interest (8, 44). (A user-defined window around a read can be used instead as a semi-experimental option.) In a second pass, similar to Piranha, peaks that have fewer reads than would be expected across the transcriptome are removed. However, a Poisson distribution is used rather than the zero-truncated negative binomial.

A different approach is used by PARalyzer for PAR-CLIP. Here, for a given position, a kernel-density-based classifier estimates a Gaussian density profile for both T-to-C mutations (signal) and also for the absence of T-to-C mutations (background). Loci where the signal is greater than the background are called as binding sites.

##### Peak width

Demarcating the width of a peak is of important biological relevance. As already noted, different types of RBPs have differing binding preferences. Some tools, such as PIPE-CLIP, cluster adjacent overlapping reads to assign peak width, but this strategy lacks biological validity as read length is more dependent on technical factors, such as RNase activity, than on RBP binding preferences.

The strategy to discern peak width from the crosslink positions usually needs to be adjusted to the RBP under study. As a result, a number of tools require the user to set this window, or clustering size, e.g. PARalyzer, Piranha, iCount. However, prior knowledge of the RBP is needed to do so effectively. In cases where it may not be available, comparing peak and motif distributions (Figures 2a and b), or RNA maps with different settings of clustering size (Figure 2d, 3b, 4) may be helpful. Our current default conditions rely on 3 nucleotide clustering windows for preliminary data exploration (Figure 2d), but crosslink sites from wider windows can be included to incorporate various types of RNA binding (Figure 3b). With this approach it is evident that PTBP1 binding at 3’ splice sites of repressed exons is highly clustered, and thus the sensitivity at this position remains the same as for raw data, while sensitivity at control exons drops (comparing Figure 3a and 3b). As further validation, this approach defines interaction sites with high sensitivity and specificity when using exons defined either by microarray (Figure 3b) or RNA-seq data (Figure 4).

Other methods utilise the read distribution to define the cluster boundaries on a statistical basis. wavClusteR uses a coverage-based algorithm called ‘mini-rank norm’ to identify the boundaries by evaluating all putative clusters using a rank-based approach. The CLIP Tool Kit uses a ‘valley-seeking’ algorithm. This uses user-defined thresholds based on the heights of local maxima within a cluster of peaks and the intervening ‘valley’ read coverage to delineate adjacent peaks. Finally, CLIPper uses cubic spline fitting to fit a curve to the peak, and defines the boundaries by excluding points on the curve that exceed the false discovery rate threshold. The precise margins for fitting the curve can be adjusted.

Taken together, the choice of the peak calling tool and settings for each tool can modify the sensitivity and specificity of data, thereby affecting the conclusions that are drawn (Figure 4). Thus, two principles can be used to determine the optimal approach for peak calling: settings should be tailored to the biology of the RBP under study and, when performing comparisons between data sets, the same tool and settings should be used.

#### Challenge 3. How to account for the variable RNA abundance?

The read count is not a direct measure of RBP affinity, or indeed even the importance of a binding site. It can be influenced by other factors, most notably RNA abundance. This varies from gene to gene, and so the count of CLIP cDNAs within a transcript, or within an intron, is a composite measure of both RBP binding affinity and abundance of the transcript or the intron. This is confirmed by the correlation between CLIP read counts and RNA-seq read counts (41). A negative control lacking the specific antibody (usually replaced by non-specific IgG) is usually performed as part of CLIP experiments, but due to the high stringency of the immunoprecipitation conditions in CLIP experiments, this negative control normally contains at least 100-fold less cDNAs than the specific experiments (15). Thus, if CLIP conditions are well optimised, the cDNA coverage from negative controls is too shallow to be used for correcting for RNA abundance.

To some extent, the CLIP data itself can be used to correct for the abundance of the different transcript regions. Most available data indicate that RBPs tend to crosslink quite broadly across their bound transcripts, such that in addition to the high-affinity binding sites that contain clustered crosslinking, many additional dispersed crosslink sites are present in the same transcripts, indicative of a low-affinity, ‘scanning’ mode of binding. The density of such broadly dispersed crosslinking depends on the abundance of transcript regions more than on the presence of specific binding motifs. Thus, the randomisation and permutation approach adopted by peak callers such as iCount, which uses the total number of CLIP cDNAs in each region to model the background distribution, implicitly models the variable RNA abundance between transcript regions.

In order to control for the impact of transcript abundance, additional data can be obtained in parallel with the CLIP experiment. RNA-seq data is the most commonly produced, and it has been used to normalise CLIP coverage within transcripts (41). Most peak calling algorithms do not include the ability to include RNA-seq or other independent count-based data for normalisation, but Piranha and ASpeak are two exceptions. Piranha uses it as a covariate in the zero-truncated negative binomial regression model for the counts, whereas ASpeak uses it to calculate an expression-sensitive background.

There are limitations of using RNA-seq. Most commonly, polyadenylated or total RNA-seq data is used. However, many, if not most RBPs strongly bind to pre-mRNA transcripts, especially to introns, which are not well covered by RNA-seq. In this case, it has been shown that normalising the CLIP data using NET-seq, which captures nascent elongating transcripts including pre-mRNA, improves recovery of binding motifs (45). An alternative approach is the generation of input libraries without immunoprecipitation (46). Here, the total lysate after treatment with RNase is loaded on the gel and transferred to the membrane. The RNAs that crosslink to all RBPs present in a selected section of the membrane are isolated and their cDNA library prepared in the same way as for the specific immunoprecipitated RBP. A similar approach has been employed for analysis of eCLIP data, where an enrichment score is calculated by dividing the cDNA count of a specific RBP at given site by the size-matched input (SMI) read count (17).

Furthermore, it is not sufficient just to consider read counts per transcript for data normalisation. The distribution of the reads along a transcript is also a factor. It has been observed with total RNA-seq that the abundance of reads along the long introns in the brain is variable, which results in a “saw-tooth” pattern (47); interestingly, the long introns (especially introns longer than 100 kb) are strongly enriched in genes that are specifically expressed in the brain (48). Transcription of introns longer than 100 kb is expected to take over 30 minutes, which is much longer than the time needed for any nuclear RBPs to assemble on introns - regardless of whether this binding is co-transcriptional or not. It is this long delay that leads to increased RBP binding to 5’ regions compared to 3’ regions of introns and the resulting saw-tooth pattern. Thus, it is expected that most nuclear RBPs should have the saw-tooth binding pattern on long introns expressed in the brain - the possible exceptions being the RBPs that bind introns only after splicing is completed, such as the branch point binding protein that binds to spliced intron lariats. Indeed, a study using iCLIP reported that most nuclear RBPs that bind to long introns in the brain show the “saw-tooth” pattern, including FUS, TDP-43 and U2AF2 (49). However, a study using CLIP-seq (i.e. the original CLIP method) reported that only FUS, but not TDP-43 has such pattern, which was the basis for concluding that FUS binds via a co-transcriptional deposition (50).

The difference between conclusions reached by iCLIP (49) and CLIP-seq (50) might reflect the differences in the quantitative nature of the two methods. Overlapping cDNAs that map to the same position on transcripts are much more common for TDP-43 than FUS, because the binding pattern of FUS is more broadly dispersed across introns. Due to its use of UMIs, iCLIP can quantify cDNAs that map to the same genomic locations, while the quantitative analysis of binding patterns across introns might be affected by PCR amplification artefacts in CLIP-seq. While the reasons for the observed differences remain to be further examined, it is clear that technical differences can affect the biological conclusions drawn from CLIP data, and thus data quality analyses are needed to aid its interpretation. Moreover, methods to normalise the data not only by the variable abundance of RNAs as a whole, but also by variable abundance between exons and introns, between different introns, and across long introns, are necessary to allow a more reliable interpretation of the binding profiles.

Finally, an important consideration for data analysis is that most RBPs are enriched in specific cellular compartment, where the abundance of available RNAs is likely to be different from that seen in RNA-seq or size-matched input libraries. As our appreciation of the RBP localisations in subcellular compartments grows, with techniques that fractionate the cell before performing CLIP (51, 52), it will be valuable to produce SMI data also for these compartments, thus controlling for the compartmental variations in RNA abundance.

#### Challenge 4: How to account for crosslinking biases?

It is well-established that there are inherent biases in the UV crosslinking reaction, with preferential crosslinking between certain peptides and certain nucleotides. UV-C induced crosslinking, as used in the truncation-based methods, occurs predominantly at uridines (12). Furthermore, analysis of the SMI controls from eCLIP experiments identified ten generically enriched tetramers (16). These generic motifs were enriched at cDNA starts of eCLIP and iCLIP data of multiple RBPs, indicating that they might reflect increased efficiency of crosslinking, rather than simply the presence of a few dominating RBPs in these different experiments. All the generic motifs have a high uridine content, which is consistent with uridine enrichment that is seen in iCLIP when using UV-C for crosslinking (12), but not when crosslinking is induced by a mutant RNA methylase in m5C-miCLIP (53).

The SMI control can be used to account for these biases as well as normalising for RNA abundance. However, it is not yet clear whether the normalisation process is sufficient, or whether peaks that overlap with those found in the SMI control should be subtracted. PureCLIP is one tool currently in development that uses a statistical framework to address this particular bias (54). It uses a hidden Markov model framework to incorporate experimental biases into the peak calling process. It learns crosslink motifs from the SMI control data for an experiment and incorporates this into the emission probability of the ‘crosslink’ state. In this way, regions that correspond both to peaks and to generic motifs can be excluded to reduce the sequence artefacts that might arise from crosslinking preferences. However, this approach should be applied with care, since the binding preferences of many RBPs may include the generic motifs; for example, proteins such as PTBP1 preferentially bind to UC-rich motifs, and therefore the generic motifs are more strongly enriched at crosslink sites in PTBP1 iCLIP data (16).

#### Challenge 5: How reproducible is the data?

As ever, CLIP experiments should be replicated to ensure the robustness of the data and the resultant biological conclusions. The overall reproducibility can be assessed to some extent by the correlation of number of crosslinks per peak. However, few tools explicitly leverage data across replicates in peak calling. One currently being developed, omniCLIP, aims to do so, in addition to modelling a number of confounding factors, including RNA abundance (55).

There are two ways to use replicates; the choice depends upon the quality of the experiment, and the desired balance between sensitivity and specificity. If the sensitivity of the experiment is a concern, biological replicates can be merged before peak calling to boost it. This is at a cost to the specificity of the results. To offset this to some extent, but still increase sensitivity, an alternative is to peak call on each replicate separately, to improve the signal-to-noise ratio, and then take the union of peaks from the replicates to maximise sensitivity. Corroborative data would be needed, of course, to validate any resultant findings.

However, if specificity is of greater importance, then after peak calling on each replicate separately, the intersect of peaks can be used. Early studies took this route to reduce the chance of peaks arising as an artefact of PCR duplication (10). Now that the use of UMIs is well-established, replicate analysis is no longer necessary to avoid PCR artefacts. Nevertheless, it does engender greatest confidence in the set of putative binding sites, provided that the sensitivity of each replicate is sufficient to allow reliable peak calling within both the highly and lowly abundant RNAs. The ENCODE consortium have refined this approach by using the irreproducible discovery rate (56) originally implemented for ChIP-seq data to identify reproducible peaks across replicates using a statistical threshold (17). Given the great variation in RNA abundance levels, it remains to be tested if this approach introduces any bias for the highly abundant RNAs.

### Modelling binding sites and the “false-negative” problem

Peak calling identifies putative binding sites, minimising the false positive rate of the underlying experimental data. There are many tools for examination of these results (Supplementary Table 3). Further simple analysis can reveal basic biological information about the RBP-RNA interaction: relationships with transcript regions, or gene sets and ontologies. However, to gain a fuller understanding, more complex characterisation is required. CLIP-methods have an intrinsic biological and computational limitation: they can only generate data about binding sites on expressed transcripts and in regions that are mappable. Furthermore, these data are restricted by the sensitivity of the experiment, as already discussed. This is termed the “false-negative” problem. To generalise the findings beyond the cell or tissue or biological state in which the experiment was performed, or indeed beyond the limitations imposed by the its quality, more complex characterisation is required. This starts with basic motif finding, but extends to computational modelling.

#### Sequence motif finding

The putative binding sites can be used to learn about the sequence preferences of the RBP under study. Motif finding tools, such as DREME (Discriminative Regular Expression Motif Elicitation) (57) and HOMER (Hypergeometric Optimisation of Motif Enrichment) (58), generally work by comparing a positive (bound) and negative (background) set of sequences and assessing the enrichment of motifs statistically (Fisher’s exact test for DREME, a hypergeometric test for HOMER) to generate position weight matrices.

The motif recognition domain may not be the RNA binding domain in the protein, hence on the transcript, the motif may not be at the binding site, but adjacent to it.So, for the positive sequence a pre-defined window around the putative binding site should be used. Care needs to be taken with the selection of the background sequences as this has a large influence over the statistical assessment of the enrichment. To maximise both sensitivity and specificity, an appropriate set of sequences should be chosen based on available knowledge of the RBP. This could be designed in silico (59), but it is probably more straightforward to select relevant genomic sequences informed by the dataset. For instance, if investigating an RBP involved in splicing, such as PTBP1, with the putative binding sites highlighting a preference for intronic binding just upstream of the intron-exon boundary, a suitable background would be the unbound deep intronic regions of the targeted genes.

Easier options such as shuffling the positive sequences (DREME) or generating a random sequence of nucleotides (HOMER), should be used only as a second-option. Shuffling will reduce the sensitivity for the detection of short motifs. A random sequence will reduce the specificity, as spurious motifs may be called significant as the true distribution of nucleotides in the genome is not random. The majority of these tools were designed for transcription factors and ChIP-seq data. Often, however, RNA motifs are shorter and more degenerate than their DNA counterparts. Recently, in *kp*Logo, a more customised tool has been developed to look for shorter sequence motifs, and also consider positional information (60). This may prove to be more useful for CLIP data.

Sequence motifs generated from CLIP data can be used to predict possible binding sites, in a genomic sequence of interest, using tools such as FIMO (Find Individual Motif Occurrences) (61). They can also be compared with those generated from in vitro experiments, such as RNAcompete (28) or RNA Bind-N-Seq (62, 63), to corroborate the specificity of the CLIP experiment. Motifs are known for only ~15% of RBPs (28), however, and poor experimental specificity should not be conflated with a lack of sequence specificity.

Although less well understood, it is known that structural context, in addition to sequence preference, plays a role in RBP binding preferences (64–67). This is likely one of the reasons for a lack of sequence specificity. It should therefore be considered when predicting binding sites. However, it is difficult to incorporate adequately either the complexity of RNA structure, or the interdependence between sequence and structure, into motif discovery tools, despite attempts to do so in tools such as Zagros (67), MEMERIS (66) and RNAcontext (68). Recent programs have been more successful, at least in incorporating the interdependence, by using a hidden Markov model (ssHMM) (69), but computational modelling of binding sites is ideally placed to integrate multiple related features, as discussed next.

#### Computational binding site modelling

GraphProt was the first tool to use machine learning methods to incorporate sequence and structure into the analysis of CLIP data (70). The features are encoded using a graph kernel approach, and a support vector machine is used to build the model, treated essentially as a classification task. Its utility in addressing the “false-negative” problem has been demonstrated: peaks not detected from the raw signal on account of being located in a poor mappability region were predicted using GraphProt, and furthermore, 90% have been experimentally validated (33). More advanced machine learning methods, such as deep boosting (DeBooster, (71)) have helped to derive more accurate predictions using multiple binding site features.

Ideally, in vivo experimental data elucidating RNA structure would be used as inputs to these models. Despite the great advances that have been made recently with the development of icSHAPE (72), DMS-seq (73), DMS-MaPseq (74) and structure-seq (75, 76) identifying paired or unpaired nucleotides; and of hiCLIP (21), PARIS (77), LIGR-seq (78) and SPLASH (79) identifying RNA duplexes, these data are not yet comprehensive enough to use for modelling. Hence, computational predictions, often using thermodynamic free energy minimisation, must be used instead despite their fallibility (80, 81). Although SHAPE data can be incorporated into these predictions (82), their inherent limitations should be borne in mind when interpreting RBP-RNA interaction preferences.

RNA sequence and structure is not the only variable that drives RBP-RNA interactions. Other factors, such as cooperative binding and position in the gene relative to exons and other features also play a role (83). These parameters can be included into both unsupervised and supervised models. iONMF uses orthogonality-regularised non-negative matrix factorisation to identify factors associated with RBP binding and estimate the importance of their contribution (83). Alternative machine learning methods, such as iDeep and iDeepS, which use neural networks, have slightly improved these predictions (84, 85).

## Understanding RBP-RNA interactions

### Integrative analysis of CLIP data across RBPs

As increasing numbers of CLIP datasets are produced for an ever-widening range of RBPs, analysis naturally turns to exploring the RNA interactions of a given RBP in the context of all the others. Several studies have already exploited CLIP data to identify co-regulatory interactions, such as the competition between hnRNP C and U2AF65 in controlling *Alu* exonsation (23); and the interplay between PTBP1 and MATR3 in co-regulating alternative splicing (86). Databases such as DoRiNA 2.0 (87) and POSTAR (88) have been set up to help. DoRiNA 2.0 uploads RBP binding sites as published. This places a severe limitation on the comparisons that can be meaningfully undertaken. As already demonstrated, both the use of different CLIP techniques and different CLIP peak callers, have a significant impact upon the number, location and size of binding sites that are discovered. POSTAR reanalyses all the raw data using a different peak caller for each kind of CLIP variant. Non-negative matrix factorisation can then be used to group together RBPs that bind to the same sites to explore co-operativity (83, 89). This does enable more reliable comparison across experiments, but it is best to avoid comparing RBPs for which different peak calling tools were used, or different types of CLIP methods, since this could result in differences that are of technical nature (89). Ideally, if a comparison is being undertaken using publically available data, the approach taken by POSTAR should be bolstered by first assessing whether the quality of the experiment is sufficient even to proceed with a comparison, and second by using the same peak calling procedure for all the RBPs that are part of the same comparison. An alternative approach has been to use matrix factorisation directly on the crosslink sites as input, thus combining binding site prediction with integration of data across RBPs (83).

### Integrative analysis of CLIP with orthogonal functional data

RNA binding profiles need to be integrated with orthogonal data to gain functional insight into the role of a given RBP-RNA interaction. Throughout this review, we have used RNA maps to demonstrate analytical considerations. However, they are also a powerful tool for studying the functions of these interactions, and understanding the position-dependent mechanisms behind these functions (Box 1) (90). Integration with non-sequencing data, such as analysis of RNA specificity with RNA Bind-N-Seq, or RBP subcellular localisation, can also provide new mechanistic hypotheses, such as the potential role of DHX30 in mitochondrial transcription termination (11).

## iCLIP, [4S]U-CLIP, [w]eCLIP

We are in an era of integrative genomics. Fusing insights gleaned from CLIP data of multiple RBPs with orthogonal genomic and non-genomic approaches will be the cornerstone for further studies of the RBP-RNA interaction networks. In order to avoid being misled in this unifying vision, it is crucial to attend to the minutiae of each data set. Here, we have considered the effects of the experimental choices on the sensitivity and specificity of CLIP data. Appreciating these limitations is necessary to adapt the computational analyses appropriately. We have discussed the need to examine sensitivity and specificity of data in combination in order to give credence to the biological conclusions drawn. A unique feature of CLIP (when compared with methods such as ChIP or RIP) is its capacity to experimentally assess specificity via visualisation of the purified RBP-RNA complexes, which also serves to check that RNase fragmentation conditions are appropriate. Moreover, computational quality controls can be performed by combining CLIP analysis with mechanistic (sequence motif) or functional data (RNA maps) regarding the RBP under investigation.

Several key steps can ensure robust assignment of binding sites from CLIP data. First, peak calling is performed to distinguish high occupancy sites from the more dispersed binding that is less likely to be of functional significance. Using cDNA starts in truncation-based methods to identify the crosslink position is crucial to maintain the single-nucleotide resolution in the peak calling step. Second, the peaks require normalising for RNA abundance and assessing for crosslinking bias. Further studies are needed to better understand how the precise parameters of both these aspects should be defined with due consideration to the binding characteristics of the RBP under study. Third, to help generalise the findings and address the false-negative problem, the peaks can be used as an input for computational predictive models of the binding sites. To achieve these goals, it is important that for each published experiment a well-annotated protocol is provided, so that computational biologists can examine the potential sources of technical variation in the data. Dozens of different CLIP protocols are already available, and further changes will likely continue to be introduced. To enable appropriate quality control analyses, we suggest that the submission of each CLIP data set to a public database is accompanied by a protocol file that describes how each of the 11 steps of the protocol were performed (7).

Due to the increasing amount of data across species, tissues, cell-lines and RBPs, the computational analysis of protein-RNA interactions is well-positioned to ask new questions. For example, it is still difficult to examine the different modes of RBP binding: could one distinguish “low-affinity, scanning” modes of binding from “high-affinity, anchored” binding from CLIP data? Other methods have generated large data sets on protein-protein interactions (91), protein localisation (92), in vitro binding preferences (28, 62) and function of RBPs (11). Integration of these diverse datasets is a present challenge, but will yield significant advances in our understanding of the role of protein-RNA interactions.

Finally, RBPs have been implicated in a range of diseases, from cancers to neurological conditions (93, 94). Studies of RBPs have already led to major medical advances. Understanding the interactions between the RBPs hnRNPA1/A2 and the SMN2 pre-mRNA has led to a breakthrough, FDA-approved treatment for spinal muscular atrophy using the antisense oligonucleotide, nusinersen (95, 96). Developing appropriate computational approaches hand-in-hand with further applications of CLIP to primary cells and tissues, pluripotent stem cell models, and disease model organisms will undoubtedly lead to further insights into protein-RNA interactions that could be targets for future therapies.

## Acknowledgements

We apologise in advance to all the colleagues whose research could not be discussed owing to space constraints. We would like to thank members of the Ule and Luscombe labs, in particular Igor Ruiz de los Mozos, Flora Lee and Federico Agostini for assistance with the tables and for valuable comments during the preparation of this manuscript. This work was supported by funding from the European Research Council (617837-Translate) to J.U., a Wellcome Trust Joint Investigator Award to J.U. and N.M.L. (103760/Z/14/Z), a Wellcome Trust PhD Training Fellowship for Clinicians Award to A.M.C. (110292/Z/15/Z), a University College London Grand Challenges Award to N.H. and the Francis Crick Institute, which receives its core funding from Cancer Research UK (FC001002), the UK Medical Research Council (FC001002), and the Wellcome Trust (FC001002).

## Summary points

#### A framework for CLIP data analysis

1. Optimising and visualising purification of the RBP-RNA complexes maximises specificity.
2. Most current CLIP protocols can amplify truncated cDNAs, and analysis of cDNA starts is the starting point for their analysis.
3. Unique molecular identifiers identify PCR duplicates, reducing downstream biases in the peak calling stage.
4. Peak calling should ideally be performed by evaluating the crosslink clusters. The window-size parameters for clustering need to be adapted to the RBP under study.
5. Size-matched input libraries are valuable to normalise the peaks for variable RNA abundance.
6. Motif analysis, in addition to providing mechanistic insight, provides insight on the quality and resolution of the data.
7. Computational modelling could help address the false-negative problem, and evaluate the contribution of RNA sequence, structure and other features to endogenous RNA recognition.
8. It is best to use a consistent experimental and analytical approach when integrating multiple CLIP data sets.

## Terms and definitions

CLIP:The key experimental method for exploring protein-RNA interactions using UV crosslinking and immunoprecipitation.
UMI:A unique molecular identifier (UMI) of random nucleotides that is introduced to the reverse transcription adapter to enable reads arising from PCR duplication to be collapsed.
SDS-PAGE:A technique to isolate proteins according to their molecular weight using sodium dodecyl sulphate (SDS) to denature the protein and polyacrylamide gel electrophoresis (PAGE) to separate them.
Peak calling:The computational process of identifying statistically significant binding sites from the experimental sequencing data.
RNA map:A tool for visualising the function of protein-RNA interactions by integrating orthogonal datasets.

## Supplementary Figure 1: Analysing eCLIP RNA maps using data from the same cell lines

RNA maps are used to explore the sensitivity and specificity of PTBP1 eCLIP data in both K562 and HepG2 cells from the ENCODE website. The regulated exons are defined using RNA-seq data following PTBP1 CRISPR knockout in K562 cells also from the ENCODE website where we observed greater signal compared to the shRNA followed by RNA-seq data. We identified the skipped exons detected using rMATS (101) using junction counts only and a P-value threshold of 0.05 and FDR threshold of 0.1. Repressed and enhanced exons were defined using an inclusion level difference threshold of 0.05; control exons were selected as those with a P-value >0.1, FDR > 0.1 and an inclusion level difference of < 0.001.

In a) we show the raw data. In b) we use peaks identified using iCount (using a 3 nucleotide peak calling half-window and 7 nucleotide clustering window). In c) We use the eCLIP peaks defined using the narrowPeaks algorithm and available from the ENCODE website.

The code to reproduce this figure is available at http://github.com/jernejule/clip-data-science

